# Challenges and opportunities for drug repurposing in cancers based on synthetic lethality induced by tumor suppressor gene mutations

**DOI:** 10.1101/2025.03.20.644275

**Authors:** Michael Vermeulen, Andrew W. Craig, Tomas Babak

## Abstract

Although two-thirds of cancers arise from loss-of-function mutations in tumor suppressor genes, there are few approved targeted therapies linked to these alterations. Synthetic lethality offers a promising strategy to treat such cancers by targeting vulnerabilities unique to cancer cells with these mutations. To identify clinically relevant synthetic lethal interactions, we analyzed genome-wide CRISPR/Cas9 knock-out (KO) viability screens from the Cancer Dependency Map and evaluated their clinical relevance in patient tumors through mutual exclusivity, a pattern indicative of synthetic lethality. Indeed, we found significant enrichment of mutual exclusivity for interactions involving cancer driver genes compared to non-driver mutations. To identify therapeutic opportunities, we integrated drug sensitivity data to identify inhibitors that mimic the effects of CRISPR-mediated KO. This approach revealed potential drug repurposing opportunities, including BRD2 inhibitors for bladder cancers with *ARID1A* mutations and *SIN3A*-mutated cell lines showing sensitivity to nicotinamide phosphoribosyltransferase (NAMPT) inhibitors. However, we discovered that pharmacological inhibitors often fail to phenocopy KO of matched drug targets, with only a small fraction of drugs inducing similar effects. This discrepancy reveals fundamental differences between pharmacological and genetic perturbations, emphasizing the need for approaches that directly assess the interplay of loss-of-function mutations and drug activity in cancer models.

**Author Summary:** Synthetic lethality is an emerging approach for targeting a biological dependency in cancer cells that does not harm normal cells. This strategy is particularly valuable for targeting loss-of-function mutations in tumor suppressor genes, which are more challenging to directly target. In an effort to accelerate treatments for cancer patients, we aimed to map out these dependencies and overlap them with responses to available drugs. We discovered different outcomes when a protein is targeted by a drug versus when that same target is disrupted genetically. Thus, if a drug is to be effectively repurposed as synthetic lethal agent, feasibility studies must capture drug biology, ideally by test the drug empirically in relevant cancer models. A second notable discovery is that in vitro synthetic lethal interactions involving cancer driver genes are significantly more likely to exhibit consistent patterns, such as mutual exclusivity in human tumor samples. This is important since selection of relevant cell lines is often critical in drug development to maximize potential for translation to clinical responses.

## 1 Introduction

Synthetic lethality (SL) provides a compelling framework for developing cancer therapies tailored to the specific genetic profiles of individual patient tumors. SL relies on the genetic interaction between two genes where disruption of either gene alone is tolerated, but their simultaneous disruption leads to cell death. Applying this strategy in oncology usually involves pharmacologically targeting a protein whose inhibition is lethal to cancer cells harboring a specific loss-of-function (LoF) mutation in a partnering gene, but spare healthy cells without these mutations (**S1A Fig**). The appeal of exploiting SL lies in the potential to increase treatment efficacy, minimize adverse effects and expand the range of actionable therapeutic targets, particularly tumor suppressor genes (TSGs) that currently lack effective therapies [1,2]. An exemplar of this approach is the use of PARP inhibitors, which harness SL to exploit DNA repair deficiencies in homologous recombination deficient (HRD) cancers, commonly driven by LoF alterations in *BRCA1* and *BRCA2* [3]. Generating over $3 billion annually, PARP inhibitors have become standard of care treatment for several HRD cancers, and are increasingly used in combination therapies that prove the value of exploiting the potential of SL-driven oncology.

Despite the success of PARP inhibitors, broader clinical adoption of additional SL drugs has not been as effective as predicted and is limited to a handful of clinical-stage developments, none of which have yet been approved. However, progress may have been limited by a lack of systematic SL mapping abilities in human cells. Recent high-throughput screening efforts hold promise for expanding SL treatments, allowing unbiased target exploration across the genome. Genome-wide CRISPR and RNA interference-based genetic screens, including those from the Cancer Dependency Map Project (DepMap), OnTarget, and Project DRIVE, have substantially expanded the repertoire of potential therapeutic targets.[4] Since it’s public release in 2018, DepMap has become an essential tool for hypothesis generation, facilitating the translation of numerous SL interactions into clinical trials.

Ongoing trials include targeting PRMT5 (NCT05732831; Tango Therapeutics), and MAT2A in *MTAP*-deleted cancers (NCT04794699; IDEAYA Biosciences), and WRN in MSI or mismatch repair deficient tumors (NCT05838768; Novartis). As novel actionable SL targets and corresponding therapeutic agents are identified, the potential of precision oncology advances toward increasingly meaningful clinical impact.

Numerous studies have developed methods for identifying additional SL targets within genetically defined contexts. Approaches range from univariate tests [5–8], linear models such as mixed-model regressions [9], ANOVA [10], and machine learning techniques including random forests (RF) [11–16], neural networks [17] and autoencoders [18,19]. Among these, RF models have gained prominence for their capacity to identify molecular biomarkers linked to gene essentiality and drug sensitivity in cancer cell lines. RF models capture complex, multivariate, and non-linear relationships that univariate tests and linear models often fail to detect. RFs can account for interactions between variables, offer interpretable feature importance rankings, and provide insights into complex biological systems. Additionally, they are computationally efficient and adept at integrating diverse omics data alongside confounding variables, making them especially well-suited for analyzing high-dimensional datasets, such as those produced by the DepMap project [11].

Despite methodological advances, statistical models often yield thousands of potential SL interactions, necessitating the use of additional filters to enrich for the most clinically actionable interactions. To achieve this, previous studies have leveraged biological pathway data [7,20], protein-protein interactions [9,10], drug sensitivity profiles [5,20] and clinical information from The Cancer Genome Atlas (TCGA) such as tumor gene expression [5], survival statistics [5] and mutual exclusivity [21,22]. Mutual exclusivity is a powerful readout for filtering SL interactions using patient tumor mutations [23]. The concept is based on the observation that certain genetic alterations rarely co-occur in tumors, implying negative selective pressure on their coexistence (**S1B Fig**). Mutual exclusivity is thus a clinical indicator for SL. Focusing on gene pairs or pathways that exhibit mutual exclusivity in tumors can enrich potential in vitro SL interactions that are more likely to translate into therapeutic benefit.

Recognizing that drugs can elicit complex pharmacological responses on cell biology, several initiatives have systematically screened drugs across extensive cancer cell line panels. Projects such as PRISM (Profiling Relative Inhibition Simultaneously in Mixtures) [12], GDSC (Genomics of Drug Sensitivity in Cancer) [24], and CTD^2^ (Cancer Target Discovery and Development) [5] have significantly advanced our understanding of drug response across a diverse range of cancer types. These datasets enable the modelling of relationships between molecular features and drug response [9,12,25], providing valuable insight into mechanisms of action (MoA) as well as biomarkers of drug sensitivity and resistance [26,27]. Integrating drug sensitivity profiles with SL interactions can highlight cases where pharmacological inhibition and genetic perturbation of the same gene/protein induces similar synthetic lethal effects, thus supporting the reliability of SL interactions.

In pursuit of robust pharmacogenomic interactions and potential drug repositioning opportunities, two previous studies have conducted comprehensive comparisons of drug and CRISPR viability. Gonçalves et al integrated drug sensitivity profiles from the GDSC with DepMap gene viability profiles, finding that approximately 25% of 376 targeted oncology drugs had effects consistent with CRISPR deletion of the same targets [9]. Similarly, using PRISM drug sensitivity profiles, Corsello et al. [12] showed that 15% of active targeted oncology drugs and 0.8% of active non-oncology drugs displayed sensitivity profiles resembling CRISPR KO of their intended targets. These findings suggest that, while pharmacological inhibition and genetic disabling of a gene can produce similar outcomes, it is more common that the two modes of targeting a gene product differ, which violates a fundamental assumption in SL studies. Recognizing that there are typically differences between pharmacological inhibition and genetic KO of a target is an important step in assessing the potential of drug repositioning, such as incorporating more elements of drug biology.

In this study, we evaluate the principles underlying SL interactions identified from DepMap pooled CRISPR KO screens to assess their potential as druggable opportunities, with a specific focus on uncovering novel genetically targeted applications for existing clinical grade drugs. In contrast with earlier studies, we found that only a small fraction of SL interactions identified in CRISPR KO screens are likely to be replicated in tumors, with a significantly higher probability for interactions involving driver genes. For the 6,550 drugs tested in cell viability screens, concordance with genetic KO was unexpectedly low, highlighting the need for repurposing approaches that effectively capture drug biology, such as empirical pharmacogenomic screens or additional mechanistic insights.

## 2 Results

### Constructing a driver gene-centric network for repurposing and targeting cancer vulnerabilities

#### Genetic dependencies network

Our approach aimed to identify mutation-specific vulnerabilities in cancer cell lines by stratifying frequently mutated genes based on mutation status and assessing differential fitness effects across genome-wide gene KO’s. Systematically evaluating each mutated gene against all potential KO’s enabled us to build a network of genetic dependencies central to our analyses. We employed an adaptation of the Pan-Cancer Inferred Synthetic Lethalities (PARIS) framework [13] which applies the Boruta algorithm – a Random Forest-based feature selection method designed to capture all significant predictive features. Through our modified framework, we identified mutations associated with enhanced sensitivity or resistance to gene inhibition, systematically mapping critical genetic dependencies across diverse cancer cell line models without the need to exhaustively test each gene pair. This approach enables a precise and scalable characterization of mutation-driven dependencies, enabling us to explore potential therapeutic targets and drug repurposing opportunities.

Our analysis utilized input data from DepMap (release 23Q2) which included detailed mutation profiles and gene essentiality scores obtained from CRISPR LoF (CRISPRi) screens across 1,087 cancer cell lines. Gene essentiality scores – which reflect the impact of gene KO’s on cell proliferation – served as response variables. The mutation data, including both hotspot and damaging mutations with detailed zygosity information were used as predictor variables. We performed this analysis across all cell lines with complete mutation and essentiality profiles, identifying interactions both in a pan-cancer context and within 13 specific cancer types. After adjusting for confounders including microsatellite instability and cancer type using linear regression (*FDR* < 0.1) and excluding interactions with insufficient mutation data (< 4 mutations), our final pan-cancer network contained 1,428 significant genetic interactions (details provided in **Methods**). To clarify terminology, within an SL interaction, we define the mutated driver gene in the cell line as the ‘source’ gene and the gene targeted by CRISPR KO as the ‘target’ gene. Comprehensive details of all interactions, both pan-cancer and cancer-type specific, are documented in **Table S1** and **Table S2**.

#### Identification of clinically relevant SL interactions

Since the source data was generated in cancer cell lines, which rarely recapitulate the complexity of primary tumors [28–30], we were interested in exploring parameters that predicted which SL interactions are more likely to translate into clinical benefit on the basis of tumor biology. We reasoned that genetic interaction with driver genes [31–33], which are mutated at epidemiologically overrepresented levels in patients with cancer, may be more likely to conserved since cell lines tend to be dependent on their driver mutations [34]. As these genes are also more frequently mutated in patient tumors, SL interactions could reveal treatment opportunities with significant potential for patient benefit. To evaluate whether driver gene SL interactions are more likely to be conserved in tumors, we analyzed mutual exclusivity (ME) using TCGA tumor mutation and copy number alteration data (**Fig 1**).

**Figure 1.**
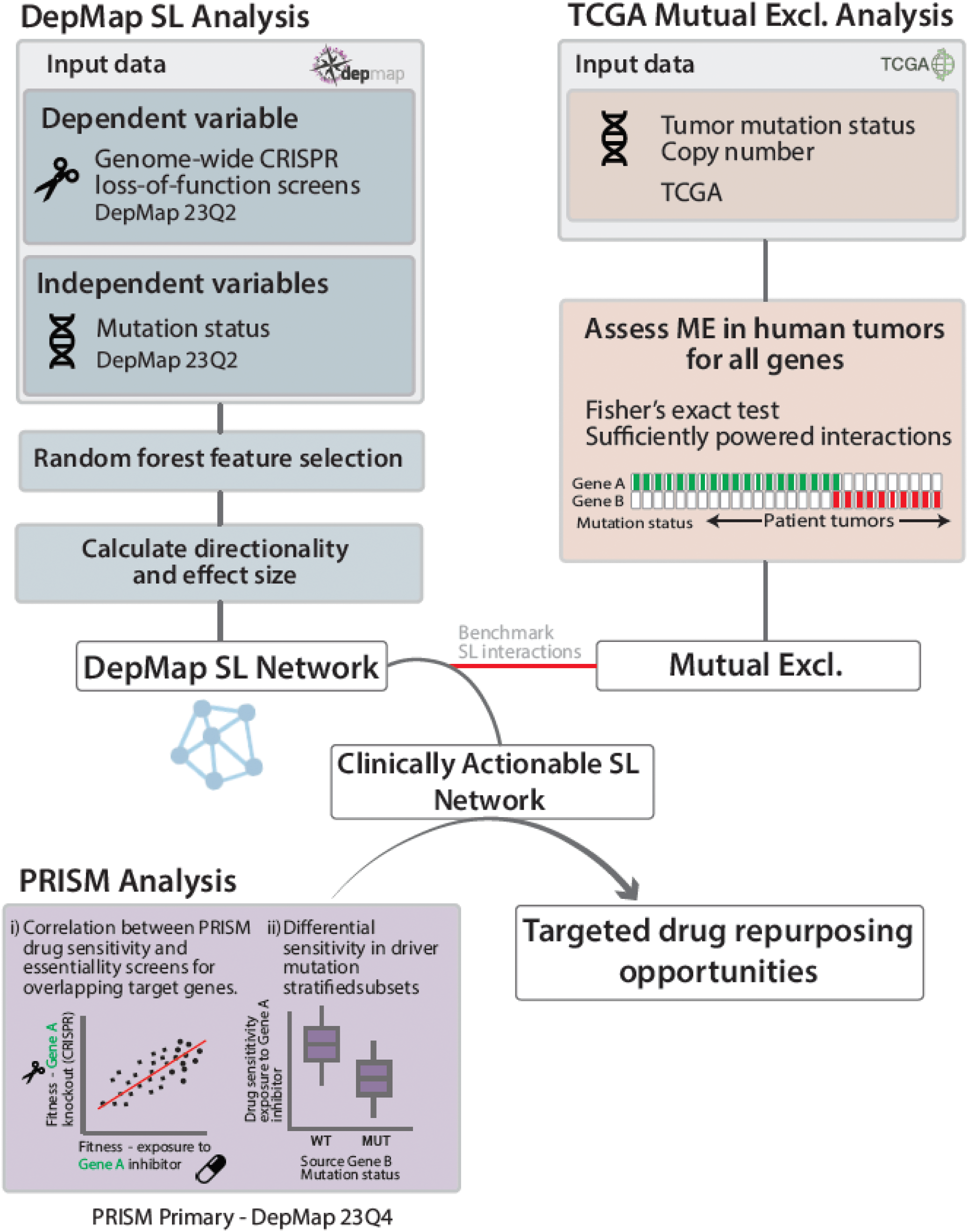
An overview of the approach to select for clinically actionable targeted drug repurposing opportunities. Using the Boruta algorithm, our pipeline integrates data from the DepMap consortium (release 23Q2) to construct a genetic interaction network. Fitness scores from CRISPR loss-of-function screens serve as dependent variables, while cell line mutation data (hotspot and damaging) are used as independent variables in a random forest (RF) feature selection process. For each perturbed gene, the algorithm assigns an importance score to mutation features, retaining those with any predictive value, and forming the foundation of our synthetic lethality (SL) network. To establish data features that are predictive of interactions holding up in the clinic (i.e. tumors), we utilized TCGA mutation data to assess mutual exclusivity (ME) across sufficiently powered genes.Non-overlapping mutations in human tumors suggest incompatibility and potential synthetic lethality. The resulting genetic interaction network was then integrated with PRISM drug screen data (release 23Q4) to identify drugs that mirror CRISPR effect scores and display differential sensitivity in driver-stratified subsets, generating a list of candidate drugs for targeted repurposing.

ME is based on the observation that certain genetic alterations co-occur less frequently than expected in patient tumors, suggesting an intrinsic incompatibility and potential SL relationship (**Fig S1B**). As a clinical indicator for SL, ME allows us to focus on gene pairs with greater translational potential. Accordingly, for each cancer type we applied hypergeometric tests to assess ME among the statistically powered gene interactions within our network (**Methods**). As hypothesized, in vitro SL interactions involving driver mutations demonstrated significantly stronger ME signals in human tumors (*P <* 0.05) compared to those lacking drivers (**Fig 2C**), even when controlling for power differences arising from higher mutation rates of drivers versus non-drivers (**Fig S2**). This observation supports the central role of driver mutations in dictating mutual exclusivity patterns and confirms that in vitro SL interactions containing drivers are biologically relevant in clinical settings.

**Figure 2.**
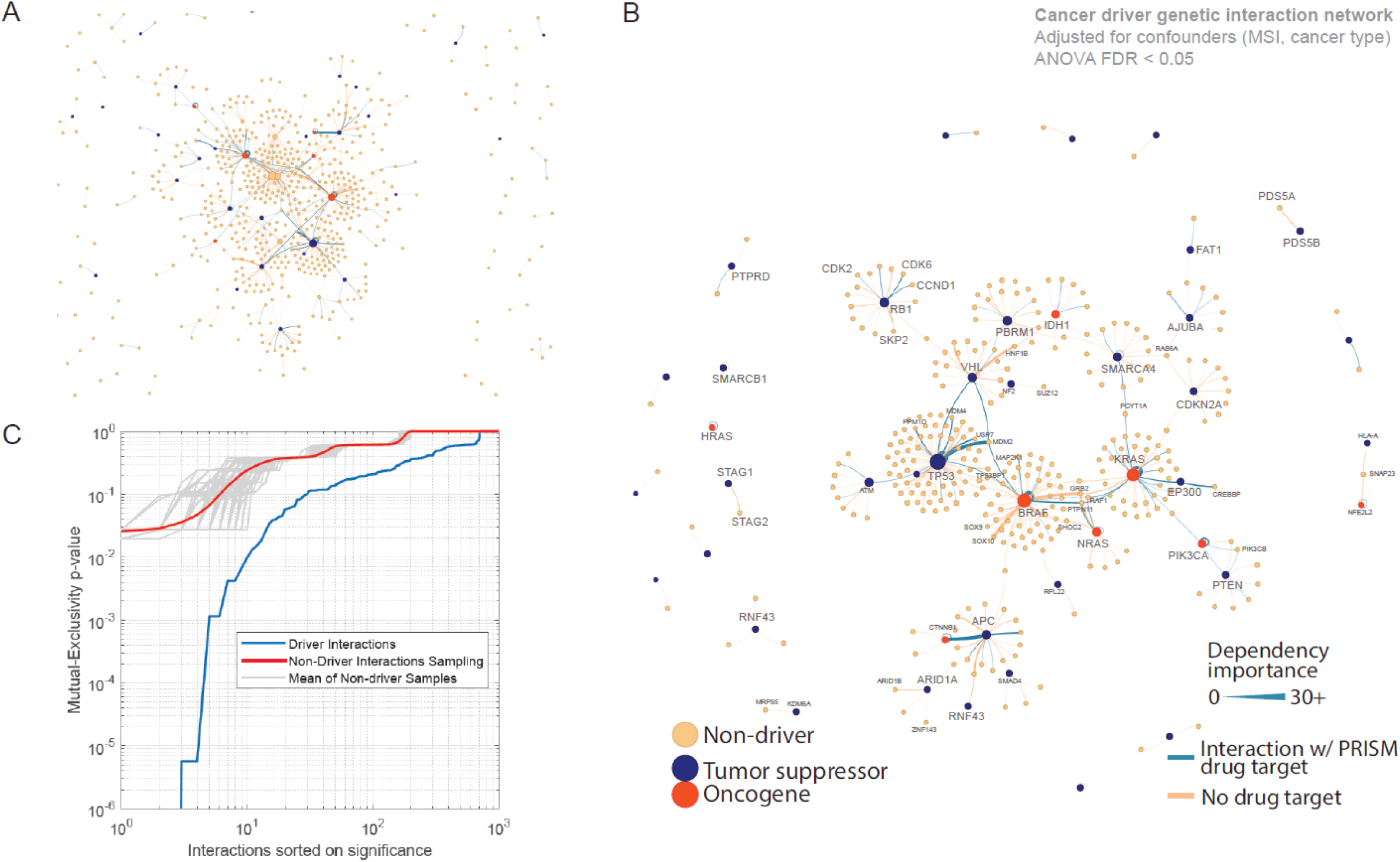
Cancer driver based genetic interaction network. A) Force-directed network of 1,428 high-confidence genetic interaction pairs identified from DepMap CRISPR viability screens, accounting for biases arising from cancer type and microsatellite instability. The network includes both synthetic lethal and synthetic viability interactions. **B)** Genetic interaction network filtered to show driver source mutations, with driver gene nodes colored red (oncogenes) and blue (tumor suppressor genes) and non-drivers in yellow. Blue edges indicate genetic interactions containing a drug target listed in the PRISM database, with edge width representing the strength of the interaction (Importance value). Node size reflects the total number of connections. **C)** Comparison of mutual exclusivity significance (y-axis) in TCGA data between synthetic lethal pairs containing a driver gene (blue) versus non-driver interactions (red) shows that interactions involving driver genes are more likely to be mutually exclusive. Grey lines represent the distribution of randomly permuted subsets of non-driver interactions.

Network visualization of the genetic interactions revealed central hubs enriched by driver genes, such as *TP53*, *APC*, *RB1* and Ras/Raf/MAPK pathway oncogenes (**Fig 2A**). Given the clustering of central hubs around common driver genes and the stronger ME signals observed in driver interactions relative to non-driver interactions, we refined the network to focus exclusively on interactions involving ‘source’ driver mutations - mutations pre-existing in the genomic profiles of the cell lines. These are interactions involving a driver mutation inherent to a cancer cell line. This subset confined the network to 510 gene pairs across 448 unique genes, including 46 driver genes (**Fig 2B**). We prioritized this subset to reduce the search space and to concentrate on interactions more likely to have clinical relevance and translational potential.

To improve interpretability, we introduced directionality to the genetic interactions using the correlation coefficient; negative values indicated increased sensitivity in the mutated state, while positive values indicated sensitivity in the wild-type. For drug repurposing purposes we were interested in SL interactions, specifically cases where mutated cell lines showed heightened sensitivity to partner gene KO.

Significant driver genetic interactions in the pan-cancer network were depicted in a heatmap (FDR < 0.1; **Fig 3A**), where numerous established dependencies were evident. These included self-paired oncogenes such as *BRAF* (Box 1) and *PIK3CA* (Box 4), paralog dependencies such as *SMARCA2-SMARCA4* (**Fig 3C**; Box 3), *STAG1-STAG2* (**Fig 3C**; Box 2), *RPL22-RP22L1*, *PDS5A-PDS5B*, and *ARID1A-ARID1B*, and other established pathway dependencies such as *SHOC2-NRAS*, *PTPN11-BRAF*, *CREBBP-EP300* and *RB1-CDK6* (Box 3). The enrichment of well-known, and experimentally confirmed genetic dependencies gave us confidence that we were capturing biologically relevant signal with power to identify clinically relevant cases downstream in our effort to repurpose cancer therapies exploiting SL interactions. Our analysis revealed extensive SL and resistance interactions (**Fig S3**).

**Figure 3.**
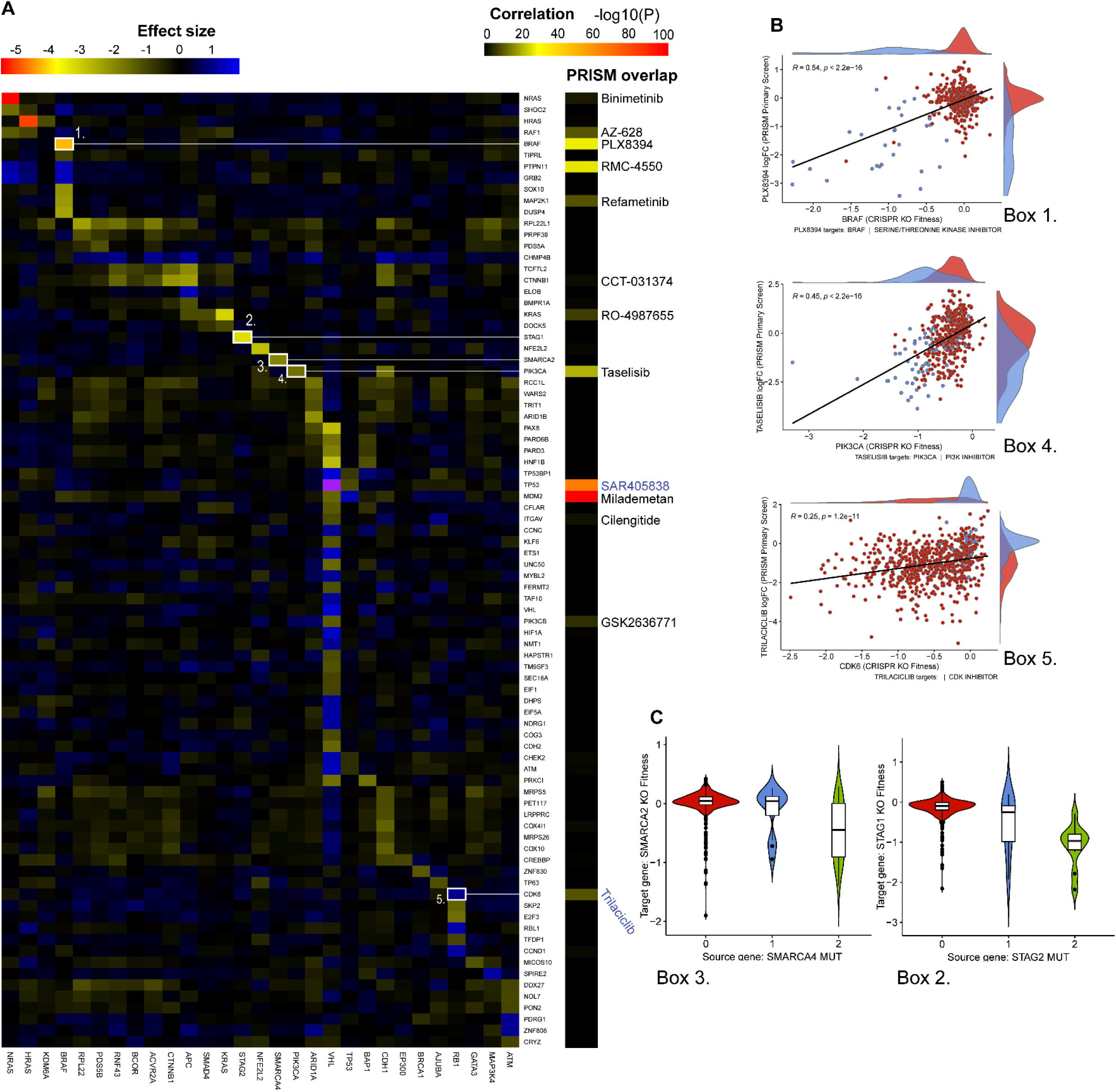
**A)** Heatmap depicting effect sizes of pan-cancer SL interactions involving driver genes, derived from DepMap CRISPR effect scores (ANOVA; *FDR* < 0.1). Driver mutations are shown on the x-axis, and CRISPR target genes on the y-axis. Negative effect size values indicate that mutated drivers sensitize cells to target gene knockout, while positive values suggest that driver mutations confer resistance or that wild-type alleles sensitize cells. On the right, Pearson correlations between PRISM drug sensitivity and CRISPR knockout effects for sample interactions are provided. Higher Pearson correlation significance indicates alignment between drug sensitivity and CRISPR knockout profiles across all CCLE cell lines, implying comparable impacts on cell viability from drug inhibition and gene knockout. Drugs with negative correlations are labeled in blue. **B)** Selected examples are highlighted in white-outlined boxes: Box 1, BRAF; Box 4, PIK3CA; Box 5, CDK6/RB1. **C)** Known SL interactions, including paralog pairs, are provided as positive controls: Box 3, SMARCA2/4; Box 2, STAG1/2.

#### Overlap with PRISM

After establishing our driver-focused genetic interaction network, we integrated 6,550 drug sensitivity profiles (6,337 unique drugs) from the PRISM 23Q4 dataset, encompassing two distinct screening efforts: Repurposing-Primary (*n* = 5362) and recently released Repurposing-1M (*n* = 1278). Drug-gene associations were mapped based on putative gene targets derived from Citeline and PRISM drug target databases. Among the pan-cancer interactions we established, 20.5% (104/510) had at least one drug with a matching target in the PRISM dataset. To explore the behavior of drug profiles targeting genes implicated in our genetic interactions, we implemented two analytical strategies: (i) examining the linear relationship between drug sensitivity and CRISPR KO effect scores, and (ii) assessing drug sensitivities in cell lines stratified by driver mutation status (**Fig 1**). Both analyses – adjusted for microsatellite instability (MSI), cell growth rates, and cancer types – aimed to identify interaction pairs where the drug effect on cell proliferation mirrored that of gene KO. While the mutation-stratified analysis is limited by smaller sample sizes, particularly in cancer-specific subsets and for less common driver mutations, it provides direct insights into context-specific therapeutic potential with translational value. In contrast, the linear correlation approach leverages the entirety of the dataset, providing better statistical power to detect strong and generalizable dependencies.

We initially assessed correlations between drug sensitivity and gene essentiality for druggable targets in pan-cancer genetic interaction pairs. Among these targets, 40.3% (42/104) displayed a significant linear correlation between drug sensitivity and CRISPR KO (*FDR* < 0.05; **S3 Table**), indicating that these genetic dependencies may be pharmacologically targeted to induce SL. Notably, many of these ’druggable SL’ pairs involved well-characterized cancer genes, such as *MDM2, TP53, MCL1, BRAF, PTPN11, PIK3CA, EGFR, BCL2L1, CDK6*, and *ERBB2*, for which targeted therapies are already available, serving as positive controls. For example, dependencies on *MDM2* and *TP53* strongly correlated with sensitivity to various MDM2 inhibitors, underscoring the parallel effects of pharmacological inhibition and gene KO on MDM2-mediated p53 repression. Other key correlations were observed with cell cycle-related genes, such as *MCL1* dependencies with MCL1 inhibitors (AZD5991, AMG-176), *CDK6* with CDK4/6 inhibitors (Ribociclib, Palbociclib, Trilaciclib), and *BCL2L1* with BCL inhibitors (Navitoclax, ABT-737; **Fig S4**). Furthermore, dependencies in epigenetic regulators—such as *EP300* with the p300/CBP inhibitor A-485 and *BRD2* with BET inhibitors including OTX015, TEN-010, and CPI-203 – highlight the potential for therapeutic intervention in these drug-gene relationships (**Fig S4**).

However, across the entire PRISM database, correlations between drug sensitivity and CRISPR KO essentiality of drug targets were rare. Of all PRISM drugs, only 154 (3%) displayed a significant correlation with any target gene essentiality profile (*FDR* < 0.1; **Table S3**). Additionally, for each drug, we compared the correlation coefficient between drug sensitivity and effect scores for all putative target genes against a distribution of randomly permuted effect scores (**Fig 4**). In the majority of cases, observed correlations did not significantly deviate from the permuted distribution, with the exception of a skewed tail of highly correlated pairs, mostly from highly targeted oncology drugs.

**Figure 4:**
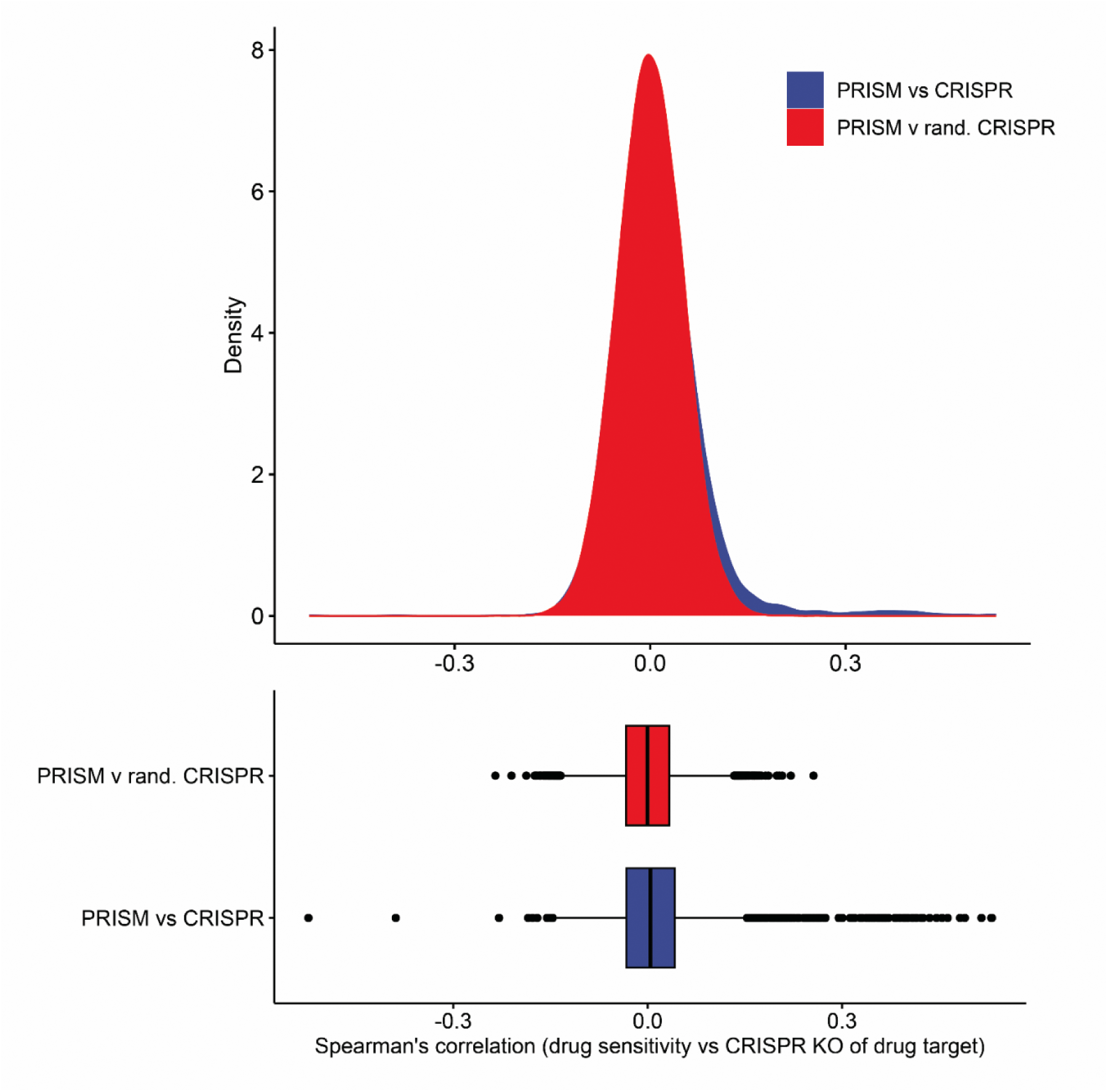
Drug perturbations typically do not replicate genetic perturbations of corresponding drug targets. Density plot showing the distribution of Spearman’s rank correlation coefficients between gene effect scores and PRISM drug sensitivity for drugs with overlapping gene targets (blue). The red curve represents the Spearman’s rank correlation coefficients between each PRISM drug and a set of randomly permuted gene effect scores, serving as a control. Bottom: Boxplot summarizing the same data, highlighting some of the strongest correlations between gene effects and drug sensitivity.

These findings suggest that while CRISPRi screens are effective at identifying essential genes, the genetic dependencies they reveal may not directly correlate with drug efficacy. Small-molecule inhibitors typically target only the catalytic function of proteins, which can activate compensatory cellular mechanisms, context-dependent dependencies, and off-target effects, complicating the translation of genetic vulnerabilities into therapeutics. Furthermore, the promiscuity of many drugs, with multiple putative targets, adds an additional layer of complexity. From a clinical translational perspective, genetically-based predictions (e.g., SL interactions) are insufficient on their own to justify drug development and rigorous validation through rigorous drug testing in model systems is essential to better validate their therapeutic potential in humans.

We then investigated whether cell lines with specific driver mutations demonstrate consistent responses to both genetic deletion and small-molecule inhibitors compared to their wild-type counterparts. We addressed this by analyzing drug sensitivities in cell lines stratified by driver mutation status, examining both hotspot oncogene mutations and deleterious mutations in TSGs across 59 driver genes (**Table S1**). Incorporating MSI as a confounder in the regression model was crucial for accurately evaluating these associations, particularly for AKT inhibitors, which demonstrated high efficacy in MSI-positive lines regardless of driver mutation status. Within our driver-focused SL network, 36 interaction pairs exhibited significant, directional agreement with drugs targeting the corresponding protein products (*FDR* > 0.2; **Table 1**). As expected, several of the strongest associations involved targeted oncology drugs in their anticipated genomic contexts—such as MDM2 inhibitors in TP53 wild-type lines, CHK inhibitors in TP53-mutated lines [35], BRAF inhibitors in BRAF-mutated lines, and CDK4/6 inhibitors in RB1 wild-type lines. Furthermore, BRAF-, NRAS-, and KRAS-mutated lines exhibited marked resistance to PTPN11/SHP2 deletion and SHP2 inhibitor RMC-4550, but not other SHP2 inhibitors. PTEN-mutated lines showed heightened sensitivity to both *PIK3CB* deletion and several pan-PI3K inhibitors. One potential novel interaction emerged, as SIN3A mutated cell lines were sensitive to nicotinamide phosphoribosyltransferase (*NAMPT*) deletion and associated NAMPT inhibitor GMX1778, although functional links are unclear.

**Table 1.** Driver-target interaction pairs showing significant directional agreement between genetic dependencies and drug sensitivities (*FDR* > 0.2). Each row lists the driver gene, target, interaction type (synthetic lethal or resistant), genetic effect, and corresponding drug effect size and targets. Color gradients indicate the direction and strength of genetic and drug effects.

| Source | Target | Driver Type | Interaction type | Genetic |  | Drug |  |  |  |
| --- | --- | --- | --- | --- | --- | --- | --- | --- | --- |
|  |  |  |  | Gini Importance | Genetic Effect | Drug Name | Drug FDR (log10) | Drug Effect Size | Drug Targets |
| TP53 | MDM2 | TSG | RESISTANT | 37.72 | 1.25 | MILADEMETAN | 35.07 | 1.13 | MDM2 |
| BRAF | BRAF | ONC | SL | 35.71 | -4.26 | PLX8394 | 18.78 | -2.91 | BRAF |
| PIK3CA | PIK3CA | ONC | SL | 11.54 | -1.7 | TASELISIB | 12.62 | -1.11 | PIK3CA |
| BRAF | MAP2K1 | ONC | SL | 5.72 | -2.3 | LIFIRAFENIB | 7.71 | -1.65 | BRAF,MAP2K1,MAP2K2 |
| RB1 | CDK6 | TSG | RESISTANT | 7.23 | 1.02 | PALBOCICLIB | 6.69 | 1.29 | CDK4,CDK6 |
| NRAS | RAF1 | ONC | SL | 3.95 | -1.36 | RAF709 | 6.54 | -1 | RAF1 |
| TP53 | CHEK2 | TSG | SL | 1.77 | -0.69 | LY2606368 | 5.16 | -0.32 | CHEK1,CHEK2 |
| NRAS | PTPN11 | ONC | RESISTANT | 5.13 | 1.13 | RMC-4550 | 3.8 | 0.79 | PTPN11 |
| KRAS | RAF1 | ONC | SL | 4.57 | -0.81 | RO-5126766 | 3.4 | -0.62 | RAF1 |
| BRAF | PTK2 | ONC | SL | 1.34 | -0.43 | CEP-37440 | 2.54 | -0.53 | ALK,PTK2 |
| BRAF | PTPN11 | ONC | RESISTANT | 10.52 | 1.27 | RMC-4550 | 1.98 | 0.98 | PTPN11 |
| KRAS | IMPDH2 | ONC | SL | 3.75 | -0.53 | MYCOPHENOLATE-MOFETIL | 1.8 | -0.16 | IMPDH1, IMPDH2 |
| TP53 | PLK1 | TSG | SL | 8.18 | -0.35 | VOLASERTIB | 1.79 | -0.3 | PLK1 |
| PTEN | MTOR | TSG | SL | 1.07 | -0.2 | NVP-BEZ235 | 1.65 | -0.6 | MTOR,PIK3CA |
| TP53 | USP1 | TSG | RESISTANT | 1.13 | 0.27 | ML-323 | 1.46 | 0.23 | USP1 |
| TP53 | ATM | TSG | SL | 2.54 | -0.69 | KU-55933 | 1.39 | -0.24 | ATM |
| PTEN | PIK3CB | TSG | SL | 0.72 | -0.62 | GSK2126458 | 1.31 | -0.54 | MTOR,PIK3CA, PIK3CB |
| ATM | PTGFR | TSG | RESISTANT | 0.23 | 0.88 | CLOPROSTENOL-(+/-) | 1.19 | 0.61 | PTGFR |
| RB1 | CDK2 | TSG | SL | 1.63 | -0.86 | PHA-793887 | 1.19 | -0.58 | pan- CDK |
| SIN3A | NAMPT | TSG | SL | 3.32 | -1.26 | GMX1778 | 1.19 | -1.08 | NAMPT |
| TP53 | PPM1D | TSG | RESISTANT | 9.99 | 0.91 | GSK2830371 | 1.19 | 0.21 | PPM1D |
| BRAF | MDM2 | ONC | SL | 6.25 | -0.82 | MILADEMETAN | 1.11 | -0.7 | MDM2 |
| BRAF | AMD1 | ONC | SL | 0.67 | -0.58 | TROMETAMOL | 1.04 | -0.17 | PTGS1,PTGS2 |
| KRAS | PTPN11 | ONC | RESISTANT | 4.81 | 0.49 | RMC-4550 | 1 | 0.28 | PTPN11 |

Next, we broadened our analysis to investigate all mutation-specific drug dependencies across the PRISM database, aiming to uncover how often these sensitivities arise beyond our predefined genetic interaction pairs. Through this expanded analysis, we identified 44 significant interactions associated with tumor suppressor gene mutations and 96 with oncogenic mutations (**Table S4**). Our findings confirmed several known vulnerabilities, such as sensitivity to the Aurora kinase inhibitor LY3295668 in SMARCA4-inactivated lines [36], and the SL of bafetinib – originally developed as a BCR-ABL and LYN tyrosine kinase inhibitor – in BRAF-mutated cell lines. Notably, the latter association is supported by recent target deconvolution studies identifying BRAF as an off-target of bafetinib [37]. This validation underscores the utility of our approach in uncovering actionable vulnerabilities across diverse driver mutations and drug combinations. Additionally, our analysis identified potentially novel drug repurposing opportunities, including the sensitivity of *KDM6A*-mutated cancer cell lines to the enoyl-acyl carrier protein reductase FABL inhibitor MUT056399 (Damaging; *FDR* = 0.00405) and the sensitivity of *FBXW7*-mutated lines to PORCN inhibitor IWP-L6 (Hotspot; *FDR* = 0.025). FABL is a key enzyme in the bacterial fatty acid biosynthesis (FAS II) pathway that has garnered interest in antibacterial drug development. However, metabolic dependencies, such as those disrupted by FABL inhibitors, are increasingly recognized in epigenetically mutated cancers, suggesting potential therapeutic relevance [38].

### Cancer type-specific analysis

We transitioned our efforts towards cancer type-specific interactions to identify context-dependent opportunities for drug repurposing, as these may better reflect the unique genetic and tumor microenvironment landscapes of different cancer types. Previous experimental knockout studies have consistently shown that SL gene pairs tend to exhibit cancer type specificity [39]. To explore these interactions, we repeated the Boruta genetic dependency analysis across 13 cancer type subsets (**Table S2**), yielding an average of 4,528 interactions per cancer type. To prioritize the highest-value interactions, we applied stringent filtering criteria, including driver mutation status, mutation sample size (n ≥ 4), and clustering based on importance scores using the heads/tails break algorithm, as previously described. This algorithm, specifically designed to address heavily tailed distributions, allowed us to isolate the most significant interactions. After filtering, 2,500 interactions remained across 13 cancer types, with 379 interactions including a druggable target (**Fig S5**).

Apart from oncogene addiction-based SL interactions involving *KRAS,* and *PIK3CA*, and *MDM2*-*TP53* interactions, most high-confidence SL interactions were rarely shared across multiple cancer types (**Fig S6**). This is likely due to context-specific dependencies, but it also reflects the challenges posed by insufficient sample sizes in some cancer types. Many cancer types lack the statistical power to identify mutation stratified dependencies in cell lines screened for both KO and drug sensitivity assays. In many cases, only a few highly mutated genes can be tested. Nevertheless, we identified some interactions that merit experimental follow-up. PBRM1-mutated renal cancer lines show resistance to *BRD2* KO and BET inhibitors. Similar findings have been made in triple-negative breast cancer models using CRISPR KO and small molecule inhibitor screens [40].

Within our bladder cancer SL network, the interaction between *ARID1A* and *BRD2* emerged as an interesting candidate (**Fig 6A**). BRD2 has been previously highlighted as a therapeutic target in *ARID1A* mutated gastric cancer [41], and ovarian clear cell carcinoma [42], suggesting it could also be of clinical interest in bladder cancer as well. The *ARID1A*-*BRD2* interaction was ranked on average 9.6 out of 47 significant SL interactions with matching drug targets in bladder cancer, combining ranks of the three importance algorithms (**Table S2**). Additionally, BRD2 knockout phenocopied BET inhibitor sensitivity in bladder cancer cell lines as BRD2 essentiality and BET inhibitor sensitivity are strongly correlated in both (*R* = 0.60; *FDR* = 8.15e-05;). In agreement with the CRISPRi interaction, ARID1A mutated bladder cancer lines were significantly more sensitive in 11 of 19 BET inhibitors – an effect that was not observed in other cancer types with recurrent ARID1A mutations (Wilcoxon; *P* < 0.05; **Fig 6B**). A significant effect was also observed in the GDSC2 dataset, where bladder cancer cell lines showed increased sensitivity to OTX015 (*P* = 0.0574; Cohen’s *d* = -1.33). When observing the interaction in more detail, BRD2 essentiality was significantly enhanced in heterozygous mutated bladder cancer cell lines (**Fig 6C**). In agreement, heterozygous ARID1A cell lines were also sensitive to several BET inhibitors available in PRISM including ARV-825 and OTX015 (**Fig 6D**). Previous studies of mouse gastric cancer in models have shown a similar dose dependent role of ARID1A where Arid1a^-/+^ tumors facilitate global loss of enhancers resulting in p53 suppression and tumor progression, whereas Arid1a^-/-^ tumors initiates p53 activation and confers a competitive disadvantage [43].

**Figure 5:**
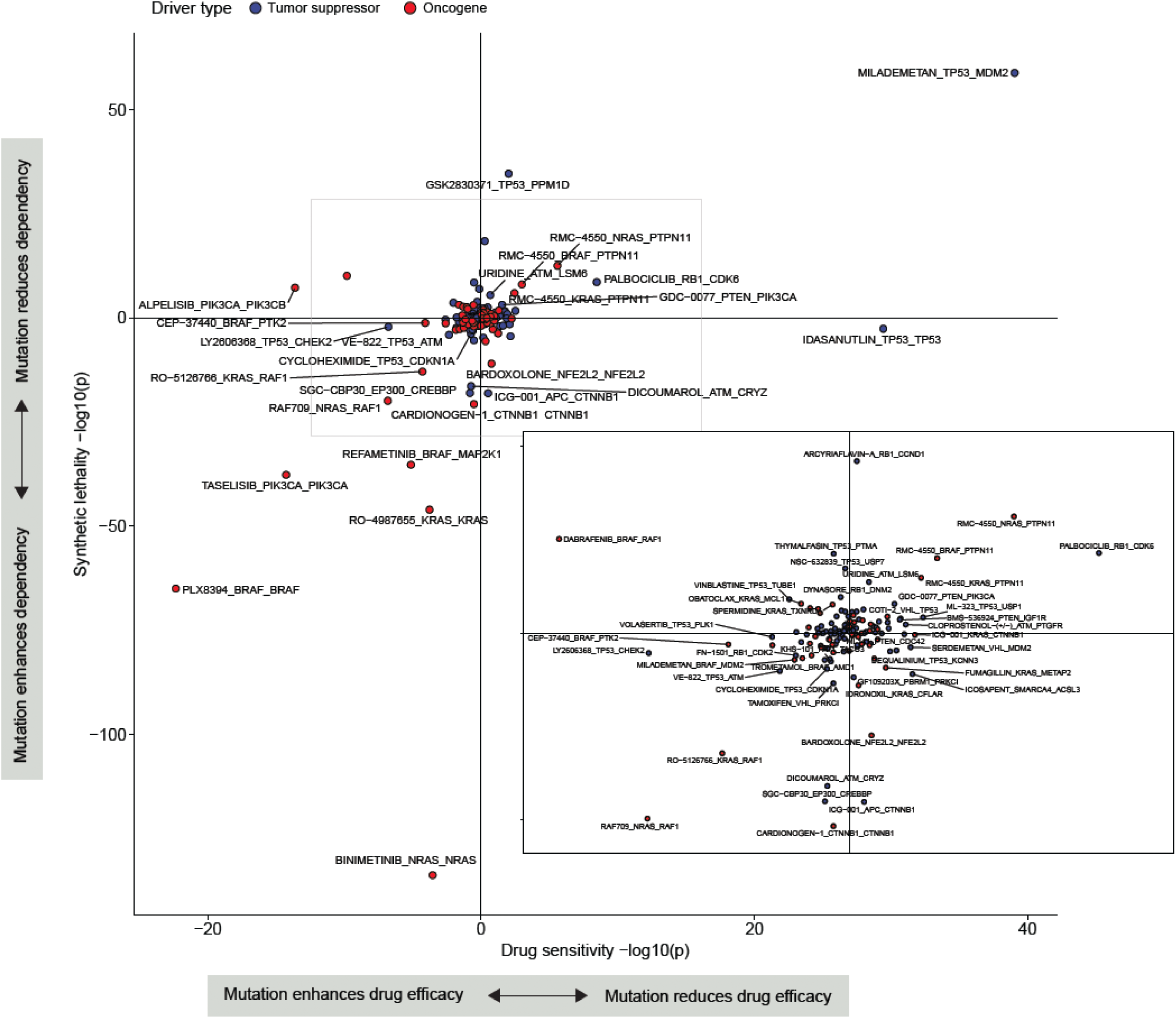
Pan-cancer synthetic lethal interactions recapitulated by PRISM drug sensitivity. Synthetic lethal interactions involving hotspot and damaging driver mutations were overlapped with PRISM drugs targeting the same genes. Tumor suppressor genes and oncogenes are indicated in blue and red, respectively. The inset provides a magnified view of the central plot region. For each SL interaction, cell lines were categorized as wild-type or mutant for the source gene, and differences between CRISPR effect scores (y-axis) and PRISM drug logFC values (x-axis) were assessed using type II ANOVA, controlling for cancer type and MSI status. Text labels follow the format “drug_source-mutation_target gene.” *P*-values were -log10 transformed and adjusted by the sign of the effect size to reflect directionality. Points in the top right quadrant have a positive effect size for both drug sensitivity and CRISPR effect, indicating decreased viability in wild-type lines for the driver gene, while points in the bottom left quadrant show increased sensitivity in driver-mutant lines to both drug and CRISPR knockout. For instance, REFAMETINIB_BRAF_MAP2K1 in the bottom left quadrant demonstrates that BRAF-mutant cell lines are sensitive to both MAP2K1 CRISPR deletion and the MAP2K1 inhibitor Refametinib. Points in the top left and bottom right suggest discordance between drug and CRISPR effects on viability. When multiple PRISM drugs matched a target gene, the most significant interaction was plotted. Full data are available in **S4 Table**.

**Figure 6.**
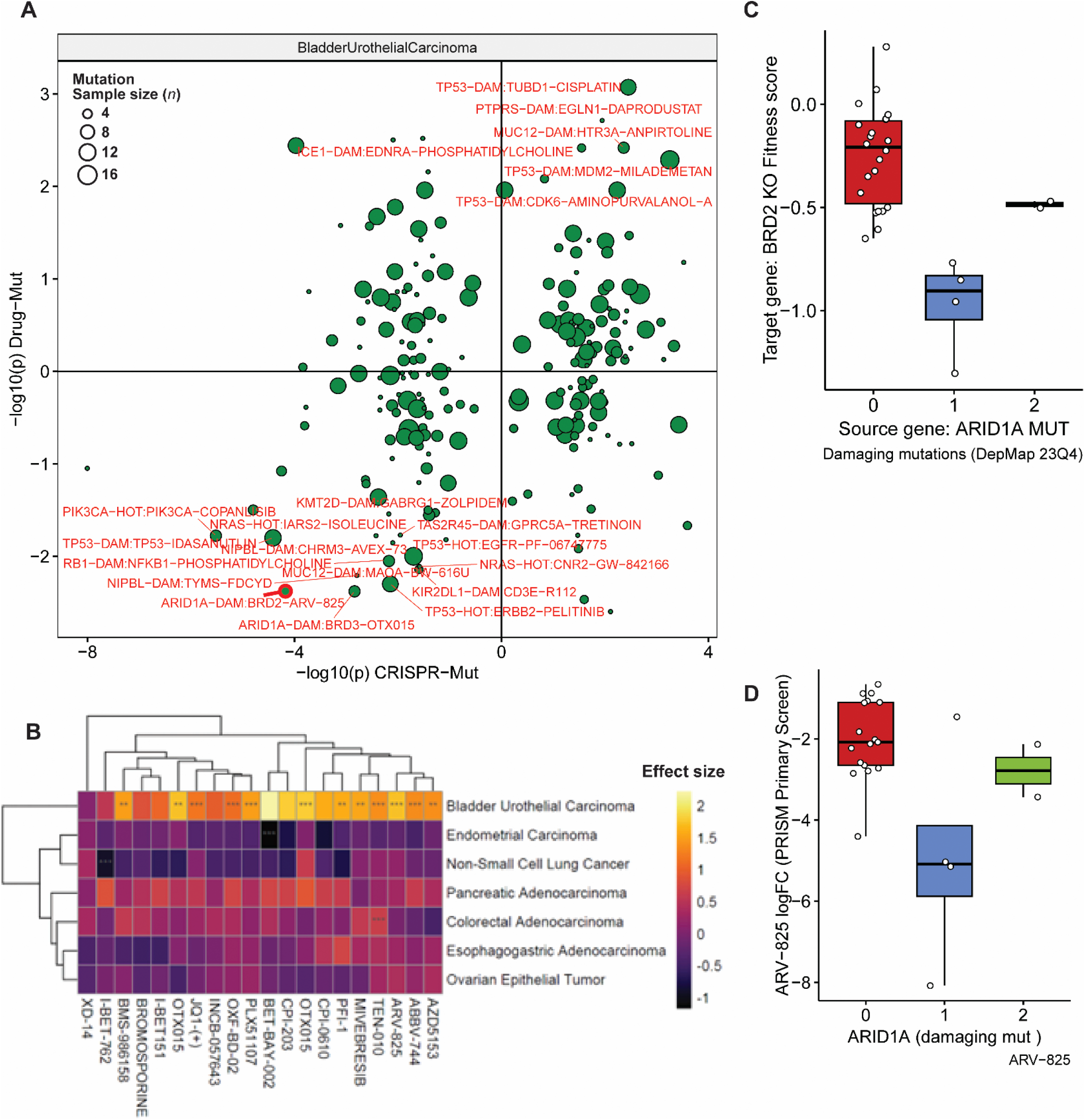
Overlap of DepMap synthetic lethal (SL) interactions and PRISM drug sensitivity in bladder urothelial carcinoma, highlighting ARID1A/BRD2 as a potential repurposing opportunity. **A**) SL interactions identified by CRISPR screens and corresponding drug sensitivities from the PRISM database in bladder urothelial carcinoma cell lines. The x-axis indicates the significance of genetic dependencies (CRISPR-Mut), and the y-axis shows the significance of drug sensitivity differences (Drug-Mut), represented as -log10(p) values. Circle size denotes sample size, while labeled red points highlight significant drug-gene interactions, notably the *ARID1A*-*BRD2* interaction with BET inhibitor ARV-825. The top gene-drug interactions are shown when multiple drugs have the same predicted targets. **B**) Heatmap depicting differential sensitivity of bladder cancer cell lines carrying *ARID1A* mutations versus wild-type cell lines to various BET inhibitors from the PRISM dataset. *ARID1A*-mutated cell lines demonstrate consistent and heightened sensitivity to nearly all BET inhibitors in bladder cancer. **C**) Boxplot showing significantly lower *BRD2* knockout (KO) fitness scores in bladder cancer cell lines harboring damaging *ARID1A* mutations compared to wild-type lines, suggesting increased dependency on *BRD2* in the heterozygous mutant context. **D**) Boxplot illustrating heightened sensitivity (negative fold change in viability scores from PRISM primary screening) of *ARID1A*-mutated bladder cancer cell lines to the BET inhibitor ARV-825 compared to wild-type cell lines.

## 3 Discussion

Despite extensive research efforts and the widespread adoption of high throughput KO and knockdown screens, PARP inhibitors (approved in 2014) remain the only available therapy leveraging SL. This leaves a significant gap between discovery and clinical application. Approximately 90% of LoF driver mutations currently lack SL-targeted therapies [44]. In this study, we systematically evaluated SL interactions involving cancer driver genes to assess their potential for drug repurposing opportunities using DepMap. Although many clinically focused, CRISPR-based, genetic SL interactions were highly statistically significant, only a subset translated effectively when mapped to drugs that target their gene product, with most successful mappings corresponding to well-characterized interactions. These findings reveal some inherent limitations of pan-cancer SL analyses and emphasize the need for drug-gene assays to improve the translatability and clinical impact of SL-based drug development.

Assuming that CRISPR-based KO of a gene fully replicates the effects of pharmacological inhibition of its encoded protein has motivated the use of CRISPR LoF screens as high-throughput proxies for identifying single-agent drugs with the potential to induce SL in cancer cells [45]. Although this approach has achieved success in certain contexts [46], significant discrepancies between the outcomes of genetic perturbation and pharmacological inhibition have been observed [47,48]. In our analysis, only 3% of drugs in the PRISM database showed a statistically significant correlation with survival effects observed in the corresponding genetic KOs, with these correlations typically confined to highly targeted oncology agents. This discrepancy is not entirely unexpected. While small-molecule inhibitors and CRISPR-mediated gene KOs can share functional similarities, they represent fundamentally different modes of perturbation. Gene deletions result in the complete loss of gene function and an abrupt cessation of expression. Although this may reveal acute cellular dependencies, it also triggers compensatory responses such as the upregulation of paralogous genes [49], adaptive rewiring of signaling pathways [50], and activation of the DNA damage response [51].

In contrast, small molecules typically target specific protein domains, often impairing catalytic activity while leaving other functional aspects of the protein, such as transcriptional and translational regulation or structural roles, intact. These drugs frequently target conserved domains shared across families of structurally related proteins. For example, BET inhibitors target bromodomains BD1 and BD2 present in BRD2, BRD3, BRD4, and BRDT [52], and Imatinib is a competitive inhibitor of ATP binding to ABL kinase and also targets the ATP-binding sites of other tyrosine kinases, including c-KIT and PDGFR [53]. As a result, inhibitors often exhibit off-target effects that complicate the specificity of their action, particularly in kinase families, where entire groups of related proteins may be inadvertently inhibited. These off-target effects not only challenge therapeutic precision but can also confound efforts to identify the true biological drivers of anti-tumor responses [54]. In some cases, off-target effects serve as the primary mechanism of tumor suppression [55], highlighting the complexity of drug responses compared to the complete LoF induced by gene KO’s.

We believe that unbiased, high throughput methods such as CRISPR KO screens remain indispensable for advancing SL drug development. However, integrating small molecules directly into genetic screens is likely crucial for capturing pharmacological effects and identifying clinically relevant SL interactions, ultimately improving the translational relevance of these findings [34].

Another major challenge in translating SL interactions identified from in vitro screens such as DepMap, as demonstrated by our work and that of others [1,11], is the pronounced context specificity of dependencies. SL dependencies can vary widely depending on factors such as cell of origin, cancer type, cellular state, and experimental conditions, including media composition during screens [56]. This variability presents significant obstacles to identifying SL interactions with broad applicability across diverse cancer types, thereby limiting their translational potential. Controlling for these contextual factors often results in diminishing returns in the discovery of novel SL targets in pan-cancer analyses, while cancer-specific analyses often suffer from insufficient statistical power. These conclusions reflect the inherent complexity of SL target discovery and highlights the need for: (i) accelerated molecular characterization, genetic and drug sensitivity screening of cell lines across diverse cancer types and, (ii) continued adoption of advanced screening strategies, such as integrating drug-gene (pharmacogenomic), gene-gene (combinatorial CRISPR), and drug-drug (combinatorial drug) KO screens, to identify robust, context-specific vulnerabilities and therapeutic opportunities.

The observed divergence between pharmacological inhibition and genetic knockout effects highlights significant limitations of current large-scale genetic screens and the challenges of using genetic perturbation to predict drug therapy effectiveness. These screens often fail to fully align drug sensitivity profiles with SL targets, emphasizing the need for a vastly expanded dependency database, alongside methodological advancements and integrative approaches to improve predictive accuracy. This aligns with the ambitious goal of comprehensively perturbing all protein-coding genes and evaluating the effects of 10,000+ drugs across 20,000+ cancer models, as envisioned by recent initiatives [57].

However, in their current state, these screening efforts are still constrained by the limited representation of many common driver genes in specific cancer contexts and the lack of experimental models for rare cancers. This highlights the need to expand and diversify experimental systems to build a more comprehensive dependency and drug sensitivity database to improve the predictive power of SL-based drug discovery.

Our study highlights key limitations in current SL target discovery approaches, including the pronounced context specificity of dependencies, the divergence between genetic KO’s and pharmacological inhibitors, and the limited representation of diverse cancer models. Addressing these challenges requires scaling up and diversifying experimental systems while leveraging advanced strategies such as isogenic screens, combinatorial CRISPR KO’s, and pharmacogenomic screens. The technology to achieve this exists, but the ambition and resources to expand these efforts at scale are now critical to facilitate the translation of in vitro discoveries into clinical breakthroughs and accelerate the development of impactful precision cancer therapies. A recent pharmacogenomics study exemplifies these challenges by conducting genetic KO screens targeting LoF drivers and evaluating their interactions with clinically approved targeted therapies and cancer drugs. Of the 32 driver-drug interactions identified, only a *KRAS* hotspot mutations with *RAF1* knockout was reproduced using DepMap data (Pearson; *P* < 0.05; **Fig. 7A**). Interestingly, *BRCA1* and *STAG2* deletions sensitized cells to PARP inhibitors in the pharmacogenomic screen; however, similar effects were absent in DepMap when stratifying cell lines by *BRCA1* and *STAG2* mutation status. This discrepancy highlights the complexity of PARP inhibitor mechanisms, which often rely on PARP trapping to stabilize PARP-DNA complexes and induce cytotoxicity. Genetic KO’s, by contrast, do not replicate the formation of these toxic drug-target intermediates or their downstream effects on genomic instability and cell death. These findings emphasize the challenges of modeling pharmacogenetic mechanisms using genetic perturbations alone and underscore the need to incorporate drug-specific effects into therapeutic development. Expanding the scale and scope of experimental systems is essential to address these gaps, improve the predictive power of SL-based drug discovery, and bridge the divide between in vitro findings and clinical applications.

**Figure 7.**
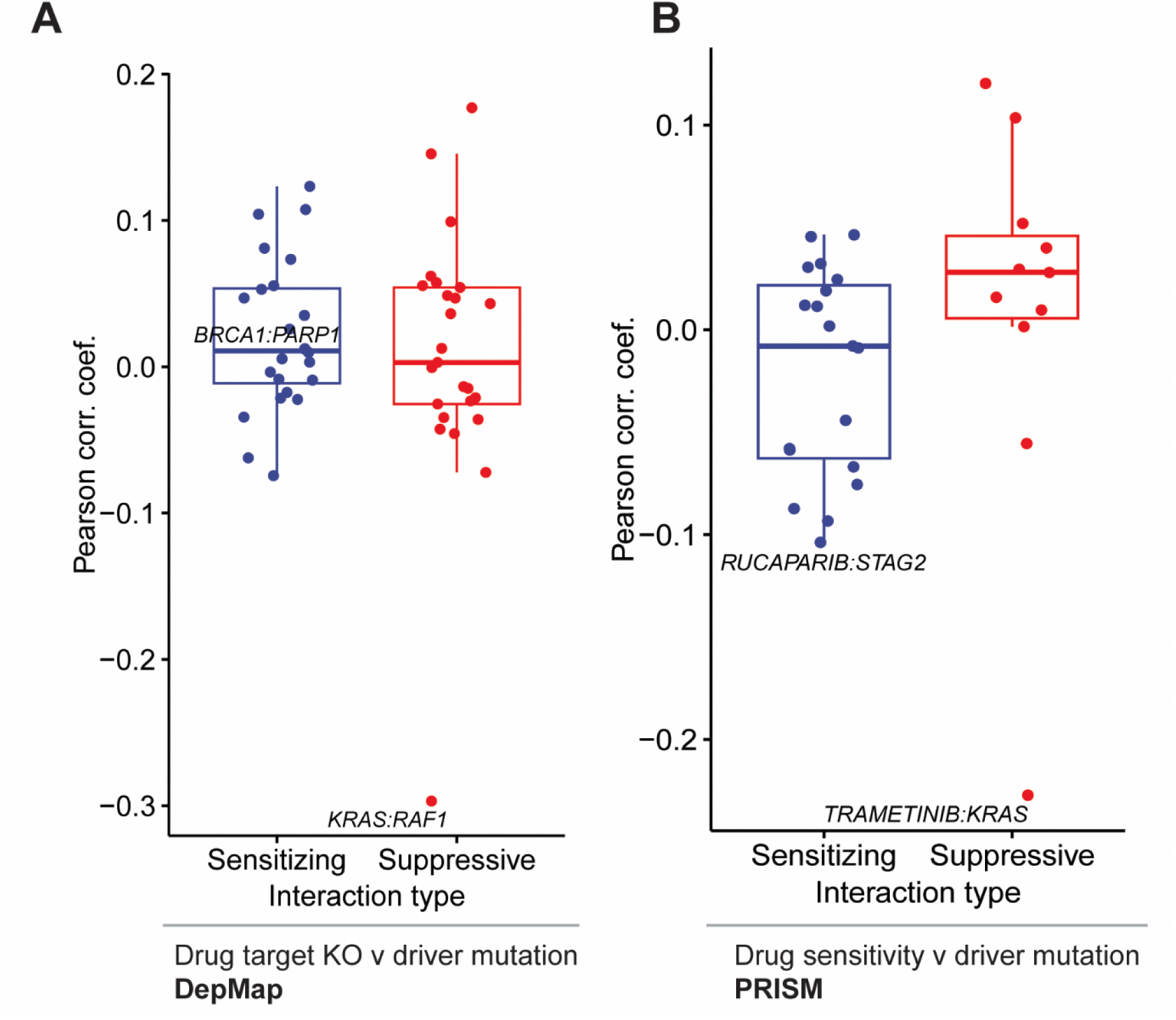
Limited replication of driver-drug interactions between pharmacogenomic and large-scale genomic screens. (A) Pearson correlation coefficients for the association between driver mutations and knockout of drug targets in DepMap CRISPR screens, stratified by interaction type (sensitizing vs. suppressive) that were detected in pharmacogenomic screens (Truesdell et al., 2024). (B) Corresponding correlations for drug sensitivity profiles from PRISM screens. Although *KRAS*–*RAF1* was reproduced in DepMap (A), most interactions, including *BRCA1*–*PARP1* and *STAG2*–RUCAPARIB, did not show consistent patterns across datasets.

## 4 Methods

### Synthetic lethal network construction

#### Input data

All cell line data, including CRISPR-Cas9 knockout fitness scores, and mutation data, were obtained from the DepMap portal (DepMap Public 23Q2; https://depmap.org/portal/). Mutation matrices were downloaded for both hotspot (OmicsSomaticMutationsMatrixHotspot.csv) and damaging mutations (OmicsSomaticMutationsMatrixDamaging.csv). As described on the DepMap portal, DepMap 23Q2 mutations were called using Mutect2, with 0 indicating no mutation, 1 for heterozygous mutations, and 2 for homozygous mutations. Hotspot mutation sites were identified based on Hess et al. (2019).

CRISPR-Cas9 knockout effect scores were extracted from the CRISPRGeneEffect.csv file. For DepMap 23Q2, gene-level knockout effect scores across all cell line models from both Achilles and Project Score were computed using Chronos and harmonized using Harmonia. Microsatellite instability scores were obtained from the DepMap Public 24Q2 release OmicsSignatures.csv file.

#### Selectively lethal genes

To refine feature selection, we excluded genes that did not significantly affect cell proliferation across all examined cell lines. Specifically, genes were retained in our dataset if they exhibited effect scores ≤ -1 or ≥ 0.4 in at least four distinct cell lines. This approach ensures that only genes influencing cell viability are included.

#### Random forest feature selection

To limit the search space and reduce the burden of multiple test correction we implemented feature selection classification using the Boruta algorithm (v8.0.0), and a workflow largely based on the PARIS pipeline (Benfatto et al., 2021). The Boruta algorithm works by first generating shadow features that are random permutations of the original features. A random forest classifier is then employed to assess the importance of each original feature relative to the maximum importance of the permuted shadow features (shadowMax). If the importance of a feature – in this case gene mutation – significantly exceeds shadowMax, it is retained; otherwise, it is rejected. In our analysis, Boruta was applied to each gene using CRISPR-Cas9 knockout effect scores as the response variable, with concatenated hotspot and damaging mutations as predictive features. Source genes with greater than 4 mutations were included, as known SL interactions were observed as significant at this threshold (**S7 Fig**). The algorithm was run across all cell lines (pan-cancer) and on cancer-specific subsets (OncotreePrimaryDisease) that had ≥ 28 fully characterized cell lines in CCLE. We employed three importance algorithms - getImpExtraGini, getImpRfZ, and getImpExtraRaw - to calculate feature importance within the random forest models (ranger v0.16.0). Interactions were considered significant only if all three algorithms concurred on feature importance. The significance threshold for feature importance was set at a p-value of 0.01, and the maximum number of iterations for Boruta was capped at 500. All code used in this study will be publicly available on GitHub: https://github.com/michaelcvermeulen/cancer_driver_repurposing (in progress)

#### Defining mutual exclusivity

Tumor mutation and copy number data was downloaded from TCGA. ME was assessed for each genetic interaction pair using Fisher’s Exact Test, a statistical method to test the non-random association between the mutation or deep deletion status of two genes across a cohort of tumor samples. For each pair, a 2×2 contingency table was constructed, with entries representing co-occurrence patterns: both genes mutated, only one gene mutated, or neither gene mutated. The resulting p-values were used to determine significant ME, with the direction of association evaluated using odds ratios: OR < 1 indicated mutual exclusivity, while OR > 1 suggested co-occurrence.

To account for cancer-type-specific effects, ME was tested separately in each TCGA cancer-type subset, with the most significant interaction retained for each interaction pair. Given the larger sample sizes associated with driver mutations, we performed random sampling of non-driver gene pairs with equal or larger sample sizes (*n*) to compare ME signal distributions. FDR correction using the Benjamini-Hochberg method was applied to adjust for multiple hypothesis testing, with gene pairs considered significant if *FDR* < 0.05.

#### ANOVA model

All pan-cancer SL interaction pairs identified through Boruta analysis were further refined to assess directionality, effect size, and potential confounding factors. Effect sizes were calculated using Cohen’s *d*, while Pearson correlations were used to establish the directionality of interactions. To account for confounding variables, including microsatellite instability (MSI), cancer type, and cell line growth properties (adherent, suspension, or mixed), we applied a Type II ANOVA model adapted from Lord et al. (2020).

The ANOVA model structure was defined as:

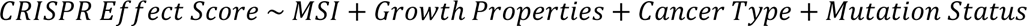

Damaging mutations were used for TSGs and hotspot mutations used for oncogenes. Mutation status *p*-values were extracted for each interaction and adjusted for multiple tests using Benjamini-Hochberg. To ensure the robustness of the final network, interactions were retained only if mutation status exhibited an *FDR* < 0.05, excluding those where confounding factors contributed significantly to the observed effects.

#### Intersection with drug sensitivity data

PRISM drug sensitivity data were obtained from the DepMap portal (DepMap Public 23Q4 release). Gene targets for each drug in the PRISM dataset were compiled by merging information from Citeline and PRISM repurposing target resources. For each pan-cancer synthetic lethal (SL) interaction, cell lines were stratified based on the mutation status of the gene of interest. Drug sensitivity, specifically to those targeting the corresponding gene, was evaluated using Cohen’s effect size, Pearson correlation coefficients, and linear models, with adjustments for microsatellite instability (MSI) status, growth properties, and cancer type. Only cell lines from cancer types with more than 10 available lines in the CCLE database, as classified by the OncotreePrimaryDisease field, were included in the analysis.

To compare the effects of targeted drugs with the CRISPR-Cas9 knockout (KO) predictions, linear models were fit between drug sensitivity profiles and CRISPR effect scores. These models were adjusted for cancer type, MSI status, and growth properties to account for confounding factors when testing pan-cancer interactions.

#### Overlap with pharmacogenomic data

Noteworthy pharmacogenomic interactions involving driver genes were sourced from Truesdell et al. [34]. Each interaction involving targeted inhibitors was evaluated in both the DepMap and PRISM datasets by assessing target gene/protein proliferative effects stratified by driver gene mutation, using both (i) the pharmacological effect of the inhibitor and (ii) the impact of genetic knockout on the corresponding target.

#### Network visualization

Pan-cancer genetic interaction networks were visualized using the visNetwork package (v2.1.2) in R, offering an interactive and customizable platform for exploring genetic interactions. Node sizes were scaled based on total connections, while edge widths were proportional to the getImpExtraGini importance scores generated by Boruta. To enhance clarity, importance scores were capped at 30. Interactions with druggable targets identified in PRISM were highlighted in blue.

## Acknowledgments

We thank Christopher D Moyes for providing valuable guidance and feedback on the manuscript. We thank Doris Coto Villa, Stephanie Young, Peter Truesdell and Ben Snider for valuable discussions and contributions to early stages of this work.

## Funding

This work was supported by a Canadian Institutes of Health Research Project Grant (PJT 178214) awarded to Andrew Craig and Tomas Babak, and funds from the Ontario Graduate Scholarship awarded to Michael Vermeulen. The funders had no role in study design, data collection and analysis, decision to publish, or preparation of the manuscript.

## Author Contributions

Conceptualization – TB, AC

Data curation – MV

Formal analysis – TB, MV

Funding acquisition – TB, AC, MV

Methodology – MV, TB

Project administration – TB, MV, AC

Supervision – TB, AC

Visualization - MV

Writing – original draft – MV, TB

Writing – review & editing – MV, TB, AC

## Supporting information captions

S1 Table. DepMap Pan-cancer SL network interactions and related PRISM drug correlations

S2 Table. DepMap Cancer-type specific SL network interactions and related PRISM drug correlations

S3 Table. Correlations between CRISPR effect score and PRISM drug sensitivity

S4 Table. Driver gene-stratified differential PRISM drug sensitivity for damaging and hotspot mutations

S1 Fig. Schematic illustrating genetic interaction concepts relevant to cancer genomics.

AS2 Fig. Distribution of synthetic lethal (SL) interaction significance values stratified by mutation frequency across cell lines.

S3 Fig. Overview of pan-cancer SL and resistance genetic interactions identified in the pan-cancer analysis.

S4 Fig. Correlation of drug sensitivity with genetic dependency profiles for MCL1 inhibitors

S5 Fig. Overview of SL interactions identified across cancer types following filtering criteria.

S6 Fig. Top cancer-specific SL interactions identified across CCLE cell lines.

S7 Fig. Relationship between the statistical significance of SL genetic interactions and the number of mutated cell lines.

## Notes

### Competing Interest Statement

T.B. is the founder and CSO of Leapfrog Bio Inc. and is the majority shareholder. All authors declare no non-financial competing interests.

## References

1. O’Neil NJ, Bailey ML, Hieter P. Synthetic lethality and cancer. Nat Rev Genet. 2017;18: 613–623. doi:10.1038/nrg.2017.47

2. Setton J, Zinda M, Riaz N, Durocher D, Zimmermann M, Koehler M, et al. Synthetic Lethality in Cancer Therapeutics: The Next Generation. Cancer Discov. 2021;11: 1626–1635. doi:10.1158/2159-8290.CD-20-1503

3. Rose M, Burgess JT, O’Byrne K, Richard DJ, Bolderson E. PARP Inhibitors: Clinical Relevance, Mechanisms of Action and Tumor Resistance. Frontiers in Cell and Developmental Biology. 2020;8. Available: https://www.frontiersin.org/articles/10.3389/fcell.2020.564601

4. Arafeh R, Shibue T, Dempster JM, Hahn WC, Vazquez F. The present and future of the Cancer Dependency Map. Nat Rev Cancer. 2024. doi:10.1038/s41568-024-00763-x

5. Srivatsa S, Montazeri H, Bianco G, Coto-Llerena M, Marinucci M, Ng CKY, et al. Discovery of synthetic lethal interactions from large-scale pan-cancer perturbation screens. Nat Commun. 2022;13: 7748. doi:10.1038/s41467-022-35378-z

6. Markowska M, Budzinska MA, Coenen-Stass A, Kang S, Kizling E, Kolmus K, et al. Synthetic lethality prediction in DNA damage repair, chromatin remodeling and the cell cycle using multi-omics data from cell lines and patients. Sci Rep. 2023;13: 7049. doi:10.1038/s41598-023-34161-4

7. Karimpour M, Totonchi M, Behmanesh M, Montazeri H. Pathway-driven analysis of synthetic lethal interactions in cancer using perturbation screens. Life Sci Alliance. 2024;7: e202302268. doi:10.26508/lsa.202302268

8. Rosenski J, Shifman S, Kaplan T. Predicting gene knockout effects from expression data. BMC Medical Genomics. 2023;16: 26. doi:10.1186/s12920-023-01446-6

9. Gonçalves E, Segura-Cabrera A, Pacini C, Picco G, Behan FM, Jaaks P, et al. Drug mechanism-of-action discovery through the integration of pharmacological and CRISPR screens. Mol Syst Biol. 2020;16: e9405. doi:10.15252/msb.20199405

10. Lord CJ, Quinn N, Ryan CJ. Integrative analysis of large-scale loss-of-function screens identifies robust cancer-associated genetic interactions. Murphy ME, editor. eLife. 2020;9: e58925. doi:10.7554/eLife.58925

11. Tsherniak A, Vazquez F, Montgomery PG, Weir BA, Kryukov G, Cowley GS, et al. Defining a Cancer Dependency Map. Cell. 2017;170: 564–576.e16. doi:10.1016/j.cell.2017.06.010

12. Corsello SM, Nagari RT, Spangler RD, Rossen J, Kocak M, Bryan JG, et al. Discovering the anticancer potential of non-oncology drugs by systematic viability profiling. Nat Cancer. 2020;1: 235–248. doi:10.1038/s43018-019-0018-6

13. Benfatto S, Serçin Ö, Dejure FR, Abdollahi A, Zenke FT, Mardin BR. Uncovering cancer vulnerabilities by machine learning prediction of synthetic lethality. Molecular Cancer. 2021;20: 111. doi:10.1186/s12943-021-01405-8

14. Dempster JM, Krill-Burger JM, McFarland JM, Warren A, Boehm JS, Vazquez F, et al. Gene expression has more power for predicting *in vitro* cancer cell vulnerabilities than genomics. Cancer Biology; 2020 Feb. doi:10.1101/2020.02.21.959627

15. Dou Y, Ren Y, Zhao X, Jin J, Xiong S, Luo L, et al. CSSLdb: Discovery of cancer-specific synthetic lethal interactions based on machine learning and statistic inference. Comput Biol Med. 2024;170: 108066. doi:10.1016/j.compbiomed.2024.108066

16. Bazaga A, Leggate D, Weisser H. Genome-wide investigation of gene-cancer associations for the prediction of novel therapeutic targets in oncology. Sci Rep. 2020;10: 10787. doi:10.1038/s41598-020-67846-1

17. Shi X, Verduzco D, Petiwala S, Jeffries C, Lu C, Anton T, et al. TCGADEPMAP – Mapping Translational Dependencies and Synthetic Lethalities within The Cancer Genome Atlas. bioRxiv; 2022. p. 2022.03.24.485544. doi:10.1101/2022.03.24.485544

18. He D, Liu Q, Wu Y, Xie L. A context-aware deconfounding autoencoder for robust prediction of personalized clinical drug response from cell-line compound screening. Nat Mach Intell. 2022;4: 879–892. doi:10.1038/s42256-022-00541-0

19. Chiu Y-C, Zheng S, Wang L-J, Iskra BS, Rao MK, Houghton PJ, et al. Predicting and characterizing a cancer dependency map of tumors with deep learning. Science Advances. 2021;7: eabh1275. doi:10.1126/sciadv.abh1275

20. Cohen Z, Petrenko E, Barisaac AS, Abu-Zhayia ER, Yanovich-Ben-Uriel C, Ayoub N, et al. SLAYER: Synthetic Lethality Analysis for Enhanced Targeted Therapy Implicates AhR inhibitor as a Target in RB1-Mutant Bladder Tumors. 2024. doi:10.1101/2024.05.01.592073

21. Fedrizzi T, Ciani Y, Lorenzin F, Cantore T, Gasperini P, Demichelis F. Fast mutual exclusivity algorithm nominates potential synthetic lethal gene pairs through brute force matrix product computations. Computational and Structural Biotechnology Journal. 2021;19: 4394–4403. doi:10.1016/j.csbj.2021.08.001

22. Liany H, Jayagopal A, Huang D, Lim JQ, Nbh NI, Jeyasekharan A, et al. ASTER: A Method to Predict Clinically Relevant Synthetic Lethal Genetic Interactions. IEEE J Biomed Health Inform. 2024;28: 1785–1796. doi:10.1109/JBHI.2024.3354776

23. Srihari S, Singla J, Wong L, Ragan MA. Inferring synthetic lethal interactions from mutual exclusivity of genetic events in cancer. Biol Direct. 2015;10: 57. doi:10.1186/s13062-015-0086-1

24. Yang W, Soares J, Greninger P, Edelman EJ, Lightfoot H, Forbes S, et al. Genomics of Drug Sensitivity in Cancer (GDSC): a resource for therapeutic biomarker discovery in cancer cells. Nucleic Acids Research. 2013;41: D955–D961. doi:10.1093/nar/gks1111

25. Carli F, Chiaro PD, Morelli M, Arora C, Bisceglia L, Rosa NDO, et al. Learning and actioning general principles of cancer cell drug sensitivity. bioRxiv; 2024. p. 2024.03.28.586783. doi:10.1101/2024.03.28.586783

26. Baek B, Jang E, Park S, Park S-H, Williams DR, Jung D-W, et al. Integrated drug response prediction models pinpoint repurposed drugs with effectiveness against rhabdomyosarcoma. PLOS ONE. 2024;19: e0295629. doi:10.1371/journal.pone.0295629

27. Pellecchia S, Viscido G, Franchini M, Gambardella G. Predicting drug response from single-cell expression profiles of tumours. BMC Medicine. 2023;21: 476. doi:10.1186/s12916-023-03182-1

28. Goodspeed A, Heiser LM, Gray JW, Costello JC. Tumor-Derived Cell Lines as Molecular Models of Cancer Pharmacogenomics. Molecular Cancer Research. 2016;14: 3–13. doi:10.1158/1541-7786.MCR-15-0189

29. Li A, Walling J, Kotliarov Y, Center A, Steed ME, Ahn SJ, et al. Genomic Changes and Gene Expression Profiles Reveal That Established Glioma Cell Lines Are Poorly Representative of Primary Human Gliomas. Molecular Cancer Research. 2008;6: 21–30. doi:10.1158/1541-7786.MCR-07-0280

30. Salvadores M, Fuster-Tormo F, Supek F. Matching cell lines with cancer type and subtype of origin via mutational, epigenomic, and transcriptomic patterns. Science Advances. 2020;6: eaba1862. doi:10.1126/sciadv.aba1862

31. Bailey MH, Tokheim C, Porta-Pardo E, Sengupta S, Bertrand D, Weerasinghe A, et al. Comprehensive Characterization of Cancer Driver Genes and Mutations. Cell. 2018;173: 371–385.e18. doi:10.1016/j.cell.2018.02.060

32. Campbell PJ, Getz G, Korbel JO, Stuart JM, Jennings JL, Stein LD, et al. Pan-cancer analysis of whole genomes. Nature. 2020;578: 82–93. doi:10.1038/s41586-020-1969-6

33. Nulsen J, Misetic H, Yau C, Ciccarelli FD. Pan-cancer detection of driver genes at the single-patient resolution. Genome Med. 2021;13: 12. doi:10.1186/s13073-021-00830-0

34. Truesdell P, Chang J, Coto Villa D, Dai M, Zhao Y, McIlwain R, et al. Pharmacogenomic discovery of genetically targeted cancer therapies optimized against clinical outcomes. npj Precis Onc. 2024;8: 1–13. doi:10.1038/s41698-024-00673-z

35. Zemanova J, Hylse O, Collakova J, Vesely P, Oltova A, Borsky M, et al. Chk1 inhibition significantly potentiates activity of nucleoside analogs in TP53-mutated B-lymphoid cells. Oncotarget. 2016;7: 62091–62106. doi:10.18632/oncotarget.11388

36. Tagal V, Wei S, Zhang W, Brekken RA, Posner BA, Peyton M, et al. SMARCA4-inactivating mutations increase sensitivity to Aurora kinase A inhibitor VX-680 in non-small cell lung cancers. Nat Commun. 2017;8: 14098. doi:10.1038/ncomms14098

37. Ji H, Lu X, Zhao S, Wang Q, Liao B, Bauer LG, et al. Target deconvolution with matrix-augmented pooling strategy reveals cell-specific drug-protein interactions. Cell Chem Biol. 2023;30: 1478–1487.e7. doi:10.1016/j.chembiol.2023.08.002

38. Xu X, Peng Q, Jiang X, Tan S, Yang Y, Yang W, et al. Metabolic reprogramming and epigenetic modifications in cancer: from the impacts and mechanisms to the treatment potential. Exp Mol Med. 2023;55: 1357–1370. doi:10.1038/s12276-023-01020-1

39. Wooller SK, Pearl LH, Pearl FMG. Identifying actionable synthetically lethal cancer gene pairs using mutual exclusivity. FEBS Lett. 2024;598: 2028–2039. doi:10.1002/1873-3468.14950

40. Shu S, Wu H-J, Ge JY, Zeid R, Harris IS, Jovanović B, et al. Synthetic Lethal and Resistance Interactions with BET Bromodomain Inhibitors in Triple-Negative Breast Cancer. Mol Cell. 2020;78: 1096–1113.e8. doi:10.1016/j.molcel.2020.04.027

41. Geng H, Qian R, Zhong Y, Tang X, Zhang X, Zhang L, et al. Leveraging synthetic lethality to uncover potential therapeutic target in gastric cancer. Cancer Gene Ther. 2024;31: 334–348. doi:10.1038/s41417-023-00706-y

42. Berns K, Caumanns JJ, Hijmans EM, Gennissen AMC, Severson TM, Evers B, et al. ARID1A mutation sensitizes most ovarian clear cell carcinomas to BET inhibitors. Oncogene. 2018;37: 4611–4625. doi:10.1038/s41388-018-0300-6

43. Loe AKH, Francis R, Seo J, Du L, Wang Y, Kim J-E, et al. Uncovering the dosage-dependent roles of Arid1a in gastric tumorigenesis for combinatorial drug therapy. Journal of Experimental Medicine. 2021;218: e20200219. doi:10.1084/jem.20200219

44. Hay M, Thomas DW, Craighead JL, Economides C, Rosenthal J. Clinical development success rates for investigational drugs. Nat Biotechnol. 2014;32: 40–51. doi:10.1038/nbt.2786

45. Fellmann C, Gowen BG, Lin P-C, Doudna JA, Corn JE. Cornerstones of CRISPR–Cas in drug discovery and therapy. Nat Rev Drug Discov. 2017;16: 89–100. doi:10.1038/nrd.2016.238

46. Xia Y, Ji X, Jang IS, Surka C, Hsu C, Wang K, et al. Genetic and pharmacological interrogation of cancer vulnerability using a multiplexed cell line screening platform. Commun Biol. 2021;4: 834. doi:10.1038/s42003-021-02352-2

47. DeWeirdt PC, Sangree AK, Hanna RE, Sanson KR, Hegde M, Strand C, et al. Genetic screens in isogenic mammalian cell lines without single cell cloning. Nat Commun. 2020;11: 752. doi:10.1038/s41467-020-14620-6

48. Lin A, Giuliano CJ, Sayles NM, Sheltzer JM. CRISPR/Cas9 mutagenesis invalidates a putative cancer dependency targeted in on-going clinical trials. Settleman J, editor. eLife. 2017;6: e24179. doi:10.7554/eLife.24179

49. Rossi A, Kontarakis Z, Gerri C, Nolte H, Hölper S, Krüger M, et al. Genetic compensation induced by deleterious mutations but not gene knockdowns. Nature. 2015;524: 230–233. doi:10.1038/nature14580

50. El-Brolosy MA, Kontarakis Z, Rossi A, Kuenne C, Günther S, Fukuda N, et al. Genetic compensation triggered by mutant mRNA degradation. Nature. 2019;568: 193–197. doi:10.1038/s41586-019-1064-z

51. Haapaniemi E, Botla S, Persson J, Schmierer B, Taipale J. CRISPR–Cas9 genome editing induces a p53-mediated DNA damage response. Nat Med. 2018;24: 927–930. doi:10.1038/s41591-018-0049-z

52. Filippakopoulos P, Qi J, Picaud S, Shen Y, Smith WB, Fedorov O, et al. Selective inhibition of BET bromodomains. Nature. 2010;468: 1067–1073. doi:10.1038/nature09504

53. Druker BJ, Talpaz M, Resta DJ, Peng B, Buchdunger E, Ford JM, et al. Efficacy and Safety of a Specific Inhibitor of the BCR-ABL Tyrosine Kinase in Chronic Myeloid Leukemia. New England Journal of Medicine. 2001;344: 1031–1037. doi:10.1056/NEJM200104053441401

54. Green MR, Newton MD, Fancher KM. Off-Target Effects of BCR-ABL and JAK2 Inhibitors. Am J Clin Oncol. 2016;39: 76–84. doi:10.1097/COC.0000000000000023

55. Lin A, Giuliano CJ, Palladino A, John KM, Abramowicz C, Yuan ML, et al. Off-target toxicity is a common mechanism of action of cancer drugs undergoing clinical trials. Sci Transl Med. 2019;11: eaaw8412. doi:10.1126/scitranslmed.aaw8412

56. Hart T, Chandrashekhar M, Aregger M, Steinhart Z, Brown KR, MacLeod G, et al. High-Resolution CRISPR Screens Reveal Fitness Genes and Genotype-Specific Cancer Liabilities. Cell. 2015;163: 1515–1526. doi:10.1016/j.cell.2015.11.015

57. Boehm JS, Garnett MJ, Adams DJ, Francies HE, Golub TR, Hahn WC, et al. Cancer research needs a better map. Nature. 2021;589: 514–516. doi:10.1038/d41586-021-00182-0

